# Assessing the impact of fire on spiders through a global comparative analysiss

**DOI:** 10.1101/2021.12.20.472855

**Authors:** Claire A. McLean, Jane Melville, Joseph Schubert, Rebecca Rose, Iliana Medina

## Abstract

In many regions fire regimes are changing due to anthropogenic factors. Understanding the responses of species to fire can help to develop predictive models and inform fire management decisions. Spiders are a diverse and ubiquitous group and can offer important insights into the impacts of fire on invertebrates and whether these depend on environmental factors, phylogenetic history, or functional traits. We conducted phylogenetic comparative analyses of data from studies investigating the impacts of fire on spiders. We investigated whether fire affects spider abundance or presence and whether ecologically relevant traits or site-specific factors influence species’ responses to fire. Although difficult to make broad generalisations about the impacts of fire due to variation in site- and fire-specific factors, we find evidence that short fire intervals may be a threat to some spiders, and that fire affects abundance and species compositions in forests relative to other vegetation types. Orb and sheet web weavers were also more likely to be absent after fire than ambush hunters, ground hunters, and other hunters suggesting functional traits may affect responses. Finally, we show that analyses of published data can be used to detect broad scale patterns and provide an alternative to traditional meta-analytical approaches.

## BACKGROUND

Fire is an important ecological and evolutionary force that has shaped ecosystems worldwide [1, 2]. While fire plays a role in maintaining species richness and ecological processes in fire-prone regions [reviewed in 3], fire can also pose a significant threat to biodiversity and individual species in sensitive landscapes or under different fire regimes [2, 4]. One of the major consequences of global climate change has been the alteration of fire regimes; in many regions, wildfires are becoming more frequent and severe, as well as threatening sensitive ecosystems that have long been fire-free [5-7]. Understanding the impacts of fire, and different fire regimes, is therefore essential for effective conservation and natural resource management now and into the future [8].

Predictive models quantifying the impacts of fire on species and ecosystems can inform fire management strategies for biodiversity conservation, including prescribed fire regimes or fire suppression. Such generalisation at broad scales is often difficult due to local variation in species assemblages and environmental factors. However, trait-based approaches, focusing on the properties of an organisms (its traits) rather than its species and relating these traits to fire responses, may overcome this challenge [9]. Additionally, an understanding of which traits predict a species’ response to fire may be useful for the management of understudied taxa. Such predictive frameworks have been used to assess the impacts of fire regimes on plant communities (e.g., [9-13]), and to a lesser extent on animals (e.g., [14-16]). One constraint of this approach is that closely related species may share similar functional traits (though not always the case; [17]), making it difficult to determine whether observed responses are due to traits or shared ancestry. However, analyses combining both trait and phylogenetic information can provide mechanistic insights into patterns of biodiversity across regions with different fire histories and help to identify species and ecosystems most at risk from changing fire regimes.

Despite their massive diversity and abundance, we have a limited understanding of the impact of fire on invertebrate communities [18, 19]. This is largely because, relative to vertebrate taxa, invertebrates have been poorly studied. It is estimated that around 80% of terrestrial arthropod species are yet to be described [20], and fundamental information about their ecology, life history, and geographic distributions remains largely unknown [21]. Invertebrates play important roles in ecosystem function and changes to invertebrate assemblages can also have knock-on effects across the food web [22, 23]. Therefore, understanding how invertebrates respond to fire is important at both the species and ecosystem level. Currently, this major knowledge gap makes it difficult to make informed fire management decisions concerning invertebrates [18, 21].

Spiders (order Araneae) are one of the most diverse and successful groups of animals [24] and it is estimated that there are approximately 80,000 species worldwide [24-26]. They are found in almost all terrestrial ecosystems, on every continent except Antarctica, and are important invertebrate predators [27]. Spiders exhibit diverse ecological, behavioural, and life history traits [28], including variation in hunting strategies, dispersal ability, and body size. The number of studies measuring the response of spiders to fire is relatively high compared to other invertebrates, likely owing to their ubiquity and high capture rates in pitfall traps (the most common survey method for invertebrates). Therefore, spiders are a potential model system as indicators of ecological disturbance [29], and can offer important insights into the impacts of fire on invertebrates and whether these depend on environmental factors, phylogenetic history, or functional traits.

Fire may impact spiders directly by increasing mortality, or indirectly by altering habitat structure and microclimate [30, 31]. A previous meta-analysis on the effects of land management (including fire) on spiders found that fire negatively affected spider abundance [32]. However, this analysis was performed for the order Araneae as a whole and more informative impacts are expected at lower taxonomic levels (e.g., family, species). Additionally, the number of studies investigating the impact of fire on spiders has grown in the past ten years. Susceptibility to fire is likely to vary across taxonomic levels due to differences in the behavioural, life-history, and morphological traits of the family or species, as well as their habitat [31]. Spiders are often classified into guilds based on prey capture methods, locomotion, and habitat use [33-35]. Such classification is useful if guild members respond in similar ways to habitat disturbances such as fire. For example, fire may have the greatest impact on taxa associated with leaf-litter, as this is the substrate most often burned during fires [19, 36, 37], while burrowing species may be sheltered from the heat and flames [38]. Additionally, the simplified habitat structure following fire may favour the more active hunting species [39], while burned habitats may be less favourable for web-building spiders if fire changes the vegetation structure and reduces the availability of web anchor sites [40, 41]. Overall, the response of animals to fire, and the adaptations and traits underpinning this, are still poorly understood [42-44].

Here, we ask the question: does fire have a global impact on spiders, and can we utilise published data to detect it? We use a phylogenetically informed framework to determine whether fire affects ground-active spider abundance or presence. We also investigate whether there is a relationship between response to fire and functional traits (hunting guild, ballooning, body length), which could contribute to the development of predictive frameworks for spiders and invertebrates more broadly. Additionally, ecosystems vary in their fire history and susceptibility, so we consider study level variation in location, vegetation type, fire type, and time of sampling in our analysis. We show that there is huge variation in spider responses to fire among studies, but that despite this there are some broad-scale patterns associated with hunting guild, vegetation type, and time since fire which may have important implications. Such information may aid land managers and policymakers to identify priority species for further study and make decisions for managing and conserving fire-prone ecosystems under a changing climate.

## METHODS

### Literature search

We conducted a systematic literature search in Web of Science on 12^th^ September 2020. Titles, abstracts, and keywords were searched using the terms “(spider* OR araneae OR arachnid* OR arthropod*) AND (*fire* OR burn*)” from any date and in any language. We also performed further unstructured searches of published and grey literature (using Google Scholar, SCOPUS, and Fire Research Institute) as well as citations within returned literature. Studies were included in our analysis if they: 1) used burned versus unburned, before versus after, or before, after, control, impact (BACI) study designs, 2) identified spiders to at least family level, 3) recorded abundance or presence/absence, and 4) used pitfall trapping. Requirement 4 was to allow us to standardise abundance measures and compare data between studies (i.e., adjust abundance according to sampling effort; more details below). Of the 1,105 studies returned by the literature search, 139 were relevant, 47 met the above criteria for inclusion in the analysis and 34 of these provided usable data with the manuscript (Table S1). There were four cases where two studies had used the same underlying data, so these were combined, resulting in 30 independent datasets (see PRISMA 2020 diagram; Figure S1). A list of the data sources used in the study is provided in the supplementary material (Table S1).

### Data collection

For each study we recorded the major vegetation type (forest, grassland, shrubland, woodland), mean time since the last fire (in months, for the fire treatment), fire type (controlled, wildfire), and latitude, and collated raw family and species level abundance and species level presence/absence data for both the fire (burned/after) and control (unburned/before) treatments. Studies with before versus after designs were only included if they sampled in the same season/s before and after the fire, as timing affects spider assemblages. For studies that sampled at multiple time points, we included data from all time points up to ten years post fire. For the few studies that compared fire frequencies (single versus multiple fires), we used data for the single fire only as many studies did not provide fire frequency information and we were therefore unable to include this as a factor in the models. Given our aim was to analyse the effects of fire per family/species (and not per study), we did not extract the overall effect sizes per study, as would be done in a traditional meta-analytical framework. Data were extracted from tables using Tabula (https://tabula.technology), or from figures using metaDigitise [45] in R (version 4.1.0; [46]). Taxonomy was checked for all datapoints and updated if necessary (i.e., we ensured the taxonomic classification of species, genera, and families were current as per the World Spider Catalog; [26]). We acknowledge that other landscape (e.g., species assemblages, fragmentation, microhabitat structure) and fire regime (e.g., frequency, severity, size) factors are likely to influence the response of spiders to fire. Testing relationships with these finer details is challenging in a global study such as this due to a lack of available data and insufficient statistical power to incorporate many factors. Here our aim was to detect broad-scale patterns which may help to identify priority species and ecosystems and guide future work at a local scale.

Sampling effort differed among studies, and there were multiple instances where experimental designs were not uniform among treatments within studies (e.g., different numbers of traps). Therefore, abundance (at the family and species level) was standardised by dividing absolute abundance by the number of plots, traps, and sampling days (sampling effort). We recognise that this is not a perfect calculation as it is based on our interpretation of the study design reported in the literature, but it was necessary to make abundance measures comparable. After calculating a standardised abundance (abundance/trap/day) per family/species per study and per treatment, we derived a measure of difference in abundance by subtracting the standardised abundance of the control treatment from that of the fire treatment, where a negative value indicated lower abundance in the fire treatment and a positive value indicated higher abundance in the fire treatment. We also classified species into one of three categories: 1) present in control, absent in fire treatment (“disappeared”), 2) present in both control and fire treatment (“no change”), 3) absent in control, present in fire treatment (“colonised”). For the family level analysis, we removed families with fewer than ten datapoints (ten fire-control differences). The presence/absence dataset was larger than the abundance dataset because not all studies measured abundance or provided enough detail for us to standardise abundance. This resulted in 564 datapoints for the family level analysis and 2,914 (presence/absence) and 2,595 (abundance) datapoints for the species level analyses.

We compiled information on hunting guild, ballooning ability, and female total body length (mm) using the World Spider Trait Database [28] and taxonomic literature listed on the World Spider Catalog [26] (Figure 1). These traits are expected to be ecologically relevant. For example, changes in vegetation structure following fire may favour some behavioural strategies over others [38-41], ballooning allows long distance dispersal and likely influences the ability of species to colonise burned areas after fire [47], and there may be a relationship between fire age and body length as size affects locomotion, dispersal ability, and desiccation resistance [16, 48, 49]. Species were classified into one of eight hunting guilds as described by Cardoso et al. [33] (ambush hunters, ground hunters, orb web weavers, other hunters, sensing web weavers, sheet web weavers, space web weavers, specialists). Given the huge diversity of spiders and the fact that many species are understudied, behavioural data were not available for all species in our study (68.3 % of the dataset). In these cases, we assigned the hunting guild of a congeneric species, or the family-level guilds of Cardoso et al. [33]. Similarly, the ballooning ability of individual species was missing for 79.9% of the dataset. Again, in cases where species data were not available, ballooning (presence/absence) was assigned based on whether it had been recorded in the genus or family [28, 47]. We must note that a limitation of this approach is that congeneric and family-level guild data may not reflect the actual hunting strategy or ballooning ability of species that lack behavioural data, as there is variation within families and genera. Body length data were from species descriptions or the World Spider Trait Database (averages were calculated if there were multiple datapoints per species). Female body length was used as there were more data available for female than male spiders, but again, we were unable to obtain body length measurements for all the species included in our study (female body length was missing for 27.7% of the dataset). Rather than perform list-wise deletion (removing datapoints with missing body length data), which can reduce statistical power and bias parameter estimates [50], we employed a data imputation procedure. The Amelia II R package [51] was first used to perform expectation-maximisation with bootstrapping for multiple imputation under an assumption of missing at random (MAR). All variables from Models 2 and 3 were included in the imputation step, and female body length was log transformed to meet the assumption of normality. We produced 10 imputed datasets (m = 10) and confirmed convergence of the chains visually. Each missing value was then replaced by the mean of its corresponding imputed values as per the single centre imputation from multiple chained equation (SICE) technique [52]. All analyses were performed with both the imputed and a reduced complete case dataset, which did not affect inferences about female body length (the imputed variable).

**Figure 1.**
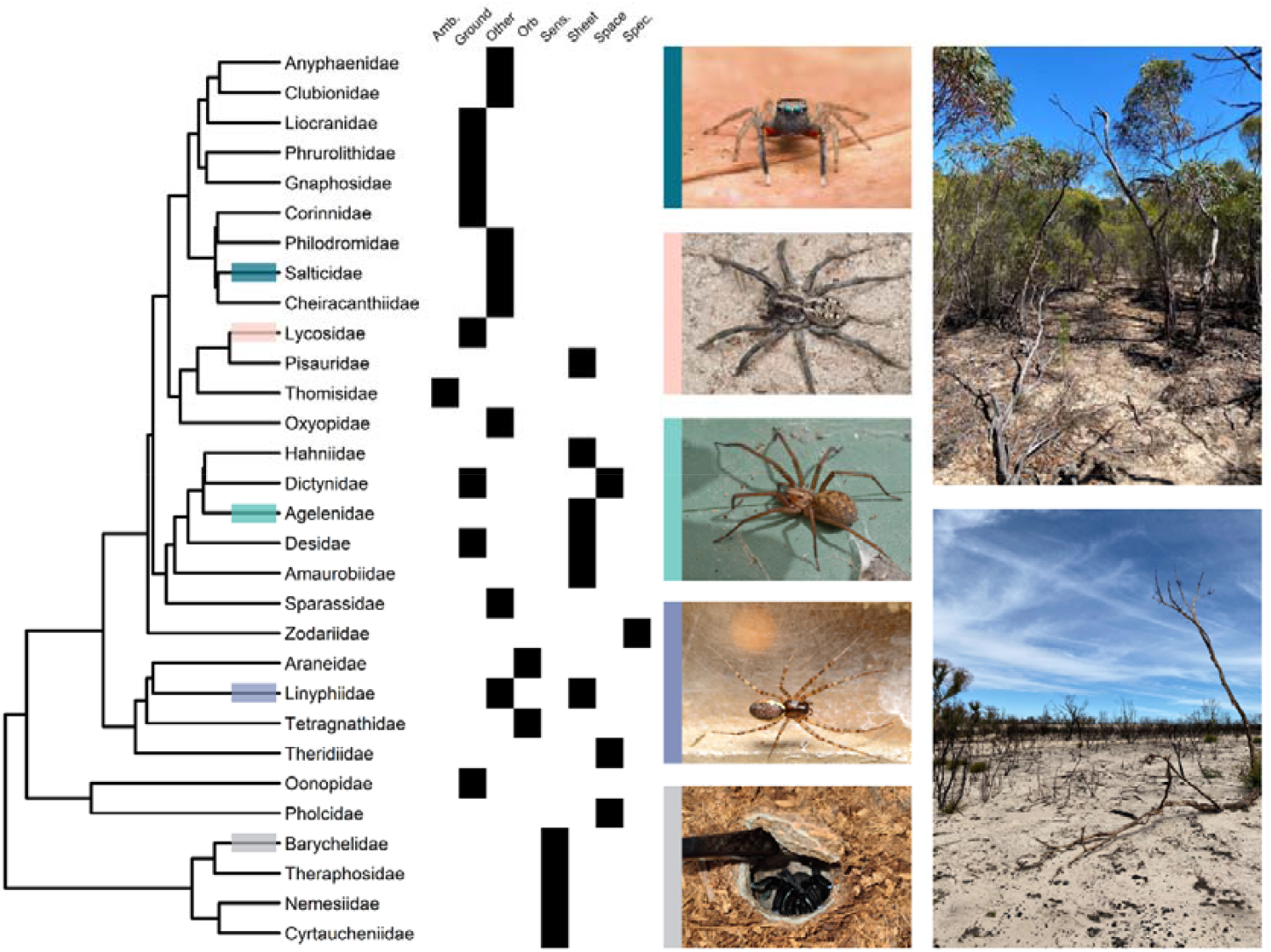
Reduced version of the spider phylogeny used in this study listing the hunting guilds present within families: ambush hunters (Amb.), ground hunters (Ground), other hunters (Other), orb web weavers (Orb), sensing web weavers (Sens.), sheet web weavers (Sheet), space web weavers (Space), and specialists (Spec.). Species from highlighted families are shown: *Jotus* sp. (Salticidae; image credit: J. Schubert), *Tasmanicosa* sp. (Lycosidae; image credit: J. Schubert), *Eratigena duellica* (Agelenidae; image credit: T. Killick), *Neriene montana* on a sheet web (Linyphiidae; image credit: T. Killick), and *Idiommata* sp. in a burrow (Barychelidae; image credit: J. Schubert). Images of unburned (top) and burned (bottom) woodland vegetation show how recent fire can alter habitat structure.

### Phylogeny and comparative analysis

Phylogenetically controlled mixed models were run using the R package MCMCglmm [53]. As a random factor, we used the phylogeny of Fernández et al. [54], which used data generated by the Spider Tree of Life project ([55]; 3 mitochondrial and 3 nuclear markers) with transcriptomic data as a backbone constraint. This was the most complete (at the family level) phylogeny available. We pruned all non-focal families using the R package phytools [56], and missing taxa were added to the tree next to sister families using the minimum branch length (1.349), in particular the infraorder Mygalomorphae was updated based on a more recent phylogeny incorporating all recognised mygalomorph families ([57]; Figure 1). For each of the models described below, we visually assessed convergence and ensured effective sample sizes were above 900. Weak priors (inverse Wishart) were used for residuals (R) and random effects (G), with variance set to 1 and different degrees of belief parameters (Table S2), and for fixed effects we used default Gaussian priors (B). Prior choice did not affect results. We extracted a statistic that quantifies the percentage of variance explained by just the fixed factors, and by both the fixed and random factors (equivalent to marginal and conditional R^2^ respectively), in each of our models [58, 59].

Using the family level dataset, we first tested whether family, vegetation type (vegetation), fire type, absolute latitude (latitude), time since fire (months), or the interaction between family and months predicted the difference in abundance between the fire and control treatment (Model 1; Table 1). We included phylogeny and the study ID as random factors to control for phylogenetic relatedness among families and multiple datapoints per study. Datapoints were weighted by the sample effort for the study (sample effort = (plots*traps*days of fire treatment + plots*traps*days of control treatment)/2) using the mev argument. We ran the model (family = gaussian) for 400,000 iterations, thinning every 100 interactions, and discarded the first 40,000 iterations as burnin. To further investigate the relationship between family and difference in abundance we ran an intercept only model (difference in abundance ∼ 1) with 3 different random factor structures, and compared them based on Akaike’s information Criterion (AIC) using the R package MuMIn [60]. The random factors used were A) phylogeny, family, and study ID, B) phylogeny and study ID, and C) study ID. This intercept-only model allowed us to explicitly quantify the separate effects of phylogenetic relationships, family, and study ID. We also calculated the phylogenetic signal (lambda) and extracted the intercept for each family from random factor model A to identify differences across families in the effect of fire.

**Table 1.**
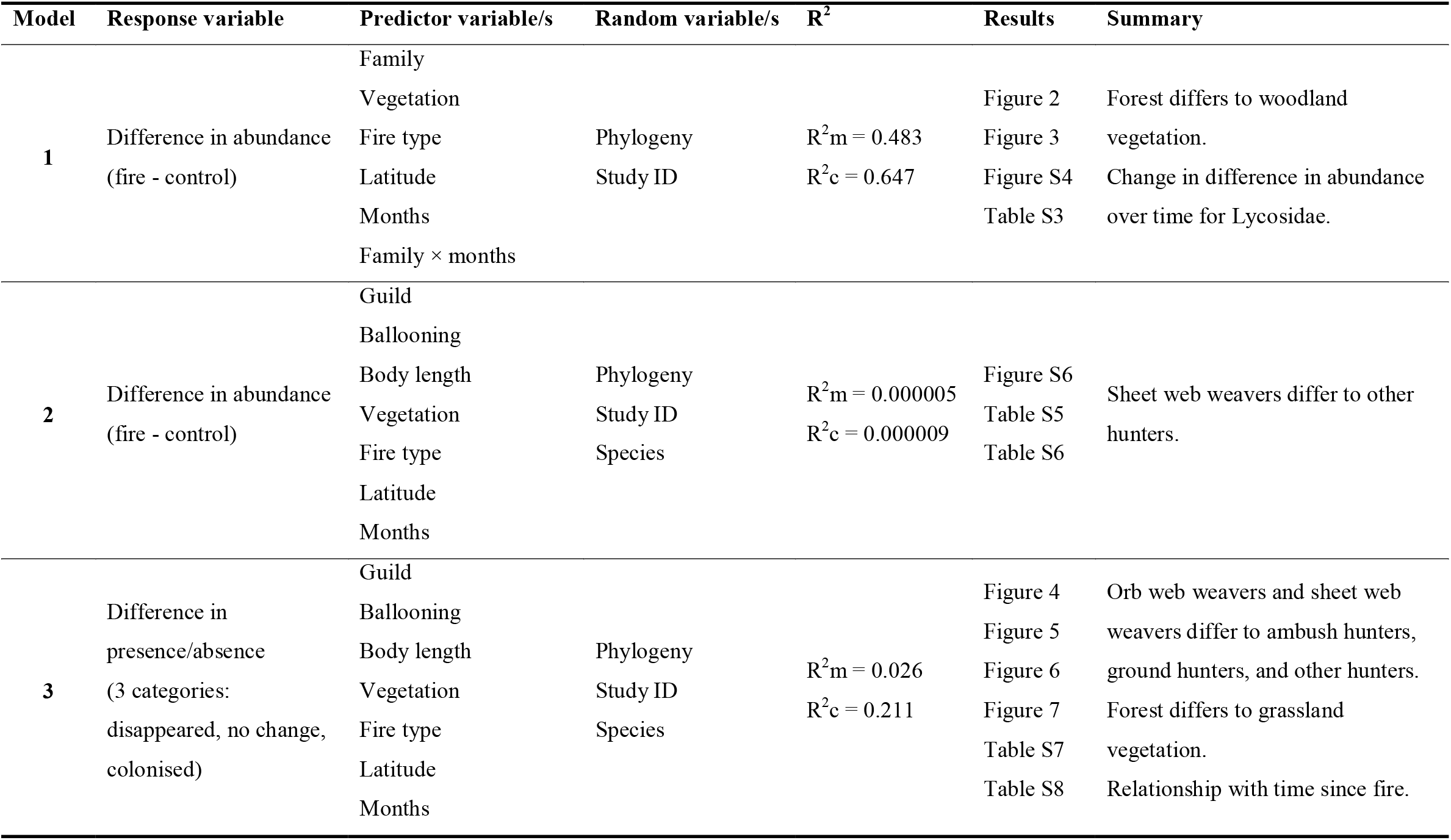
Summary of the three models tested in this study and their main findings.

We next used the species level abundance dataset to test whether hunting guild (guild), ballooning, female body length (body length), vegetation, fire type, latitude, or months predicted the difference in abundance between the fire and control treatment (Model 2; family = gaussian; Table 1). Phylogeny, species, and study ID were included as random factors, and datapoints were again weighted by sample effort (mev). Finally, we tested whether the same seven variables predicted the presence/absence of species in the control and fire treatments (Model 3, family = threshold; Table 1). Given that the response variable was an ordinal factor for this model, we fixed the residual variance to 1 (Table S2). Models 2 and 3 were both run for 150,000 iterations, thinning every 100 iterations, with the first 15,000 iterations discarded as burnin.

## RESULTS

### Summary of literature used

We analysed data from 34 studies (30 datasets; Table S1), incorporating 67 families and 1,329 species. The data spanned 16 countries across 6 continents, with North America having the most studies (11; Figure S2). The most common vegetation type was forest (16 studies), and most studies used a burned versus unburned study design (31 studies) and/or measured the effect of controlled burns (16 studies). Only a few studies provided details about fire intensity, severity, frequency, or the fire history of study sites (e.g., the age of the long-unburned site) and we were therefore unable to include this valuable information in our analysis.

### Comparative analysis

Most families showed both positive and negative differences in abundance between treatments (Figure S3). Time since fire (months) had a significant effect on the difference in abundance between fire and control treatment for the family Lycosidae (Model 1; marginal R^2^ = 0.483, conditional R^2^ = 0.648; Table 1). The relationship was positive with abundance initially lower in the fire treatment, but higher in the fire treatment from around two years after fire onwards (Figure 2). This relationship was significantly different to that observed in multiple other families (Table S3). Additionally, forest vegetation differed from woodland vegetation, and there was a marginally non-significant (P = 0.067) difference between forest and grassland vegetation. The mean difference in abundance was positive (higher abundance in the fire treatment) for forest, and negative (lower abundance in the fire treatment) for all other vegetation types (Figure 3).

**Figure 2.**
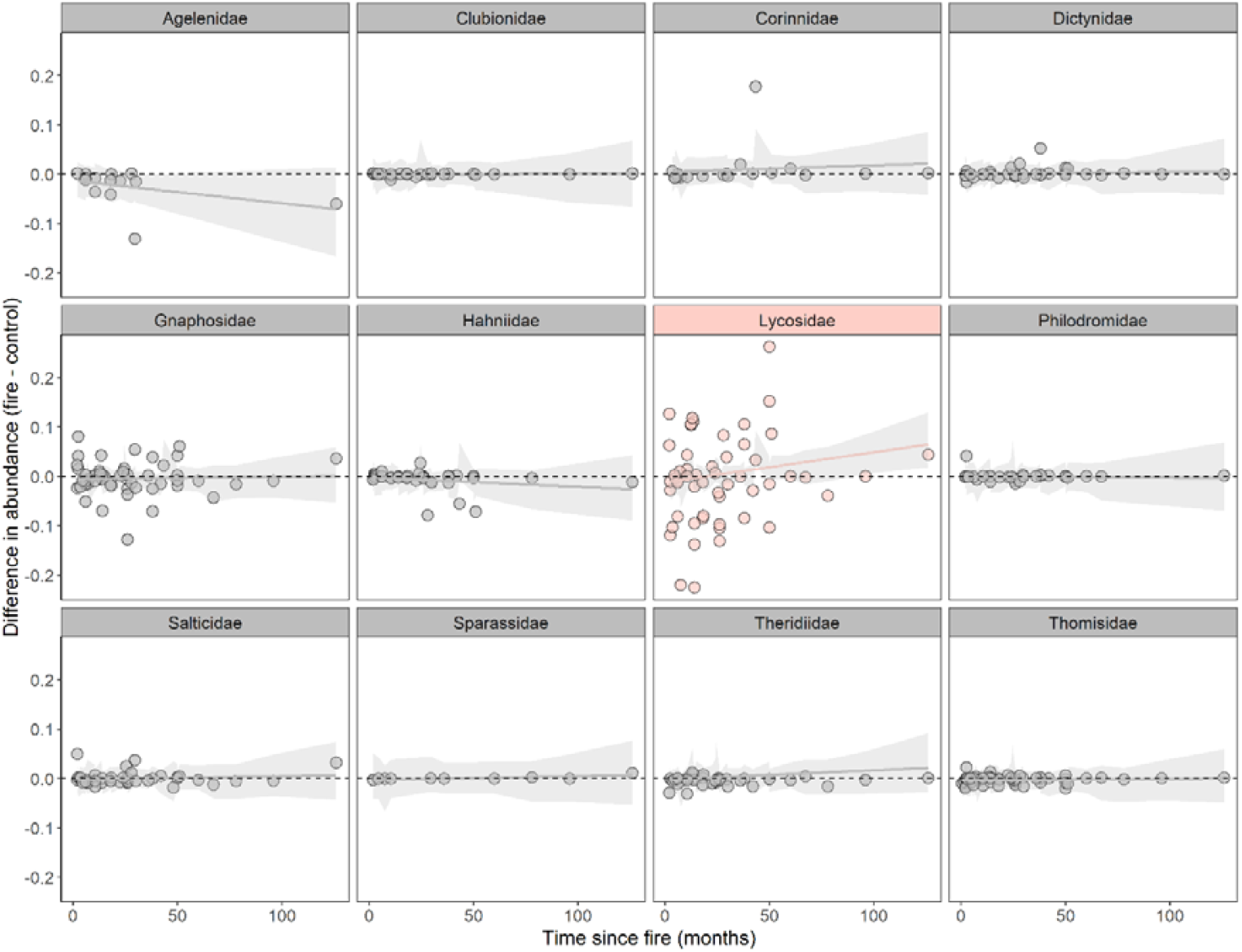
Relationship between time since fire (months) and difference in abundance between fire and control treatments within each family (Model 1). There was a significant effect of months for the family Lycosidae, and this relationship differed to all other families shown here (Table S3). Three outlying datapoints were removed for visualisation. Grey shading = 95% confidence bounds (N = 564).

**Figure 3.**
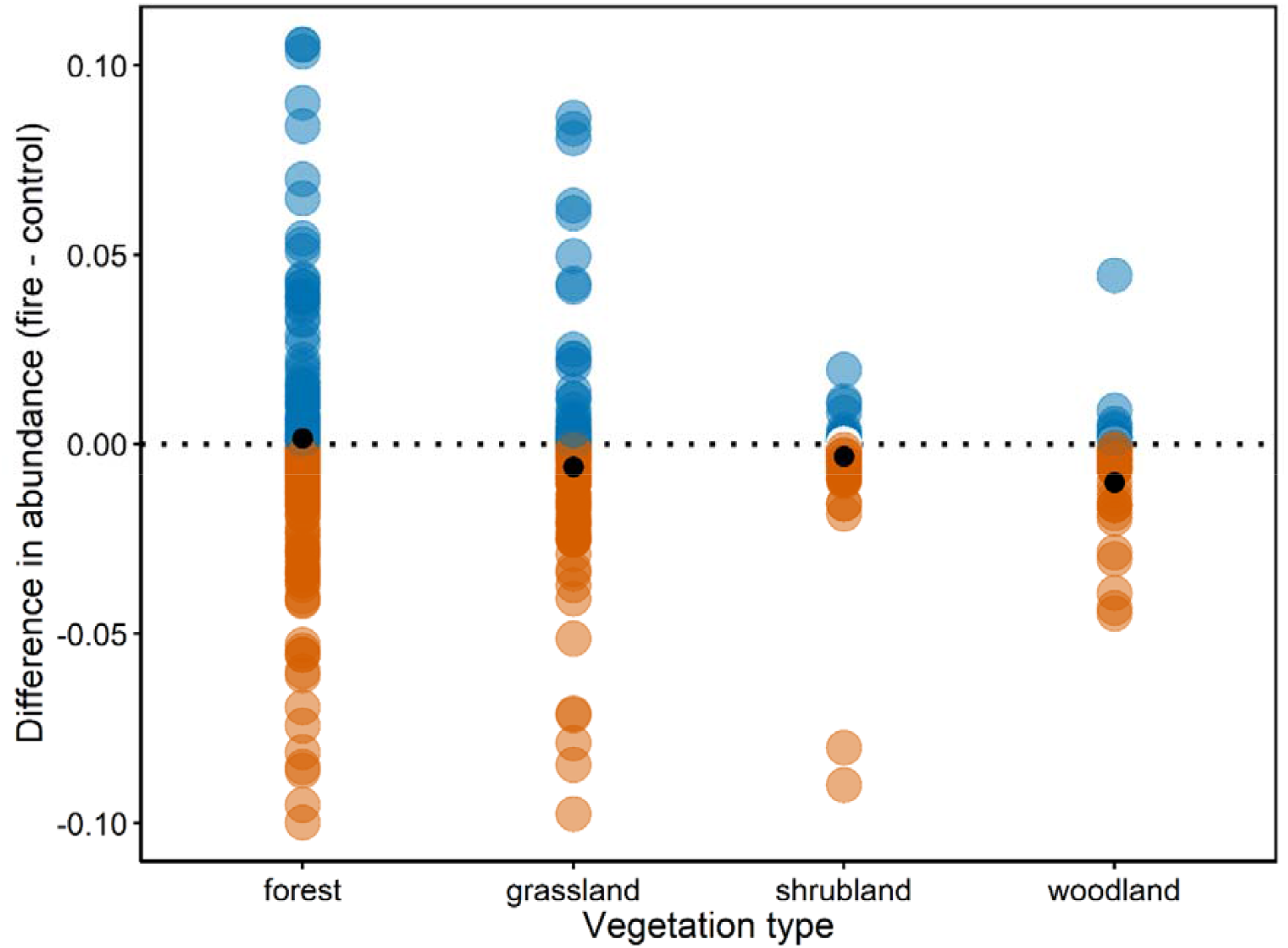
Difference in abundance between fire and control treatments for each vegetation type, for each family in each study. Blue = higher abundance in fire treatment, red = lower abundance in fire treatment, white = no difference, small black dot = mean difference in abundance for that vegetation type. There was a significant difference between forest and woodland vegetation types. Plot has been zoomed in to abundance differences between -0.1 and 0.1 for visualisation (N = 564).

Phylogenetic relatedness among families was not a strong predictor of the difference in abundance between fire and control treatments, as confirmed by examination of the random effect of family (Figure S5; Table S4), and the posterior probability of the phylogenetic signal (lambda = 0.016, 95% CI = 1.15 × 10^−7^ – 0.057) in the intercept only model. Abundance tended to be lower in the fire treatment for the families Agelenidae and Anyphaenidae (in > 80% of studies including these families there was lower abundance in the fire treatment and mean differences in abundance < 0; Figure S3), although these families had a relatively low number of datapoints, so we interpret this with caution (14 and 12 fire-control differences respectively). Conversely, multiple families with a high number of datapoints (≥30) showed mean differences of close to zero (Figure S3) and no change in abundance over time (e.g., Gnaphosidae, Hahniidae, Salticidae, Theridiidae, Thomisidae; Figure 2). Therefore, the effect of fire on each family likely differed between studies and could be either positive or negative.

Guild, ballooning, body length, vegetation, fire type, latitude, and months explained only a small proportion of the variance in the difference in abundance between fire and control treatments (Model 2; full model marginal R^2^ = 0.0000046, conditional R^2^ = 0.0000087; Table 1). Sheet web weavers differed from other hunters, and there was a non-significant trend between sheet web weavers and both ambush hunters and ground hunters (P = 0.59; Figure 4), and between forest and grassland vegetation types (P = 0.086; Figure S6; Table S5; Table S6).

**Figure 4.**
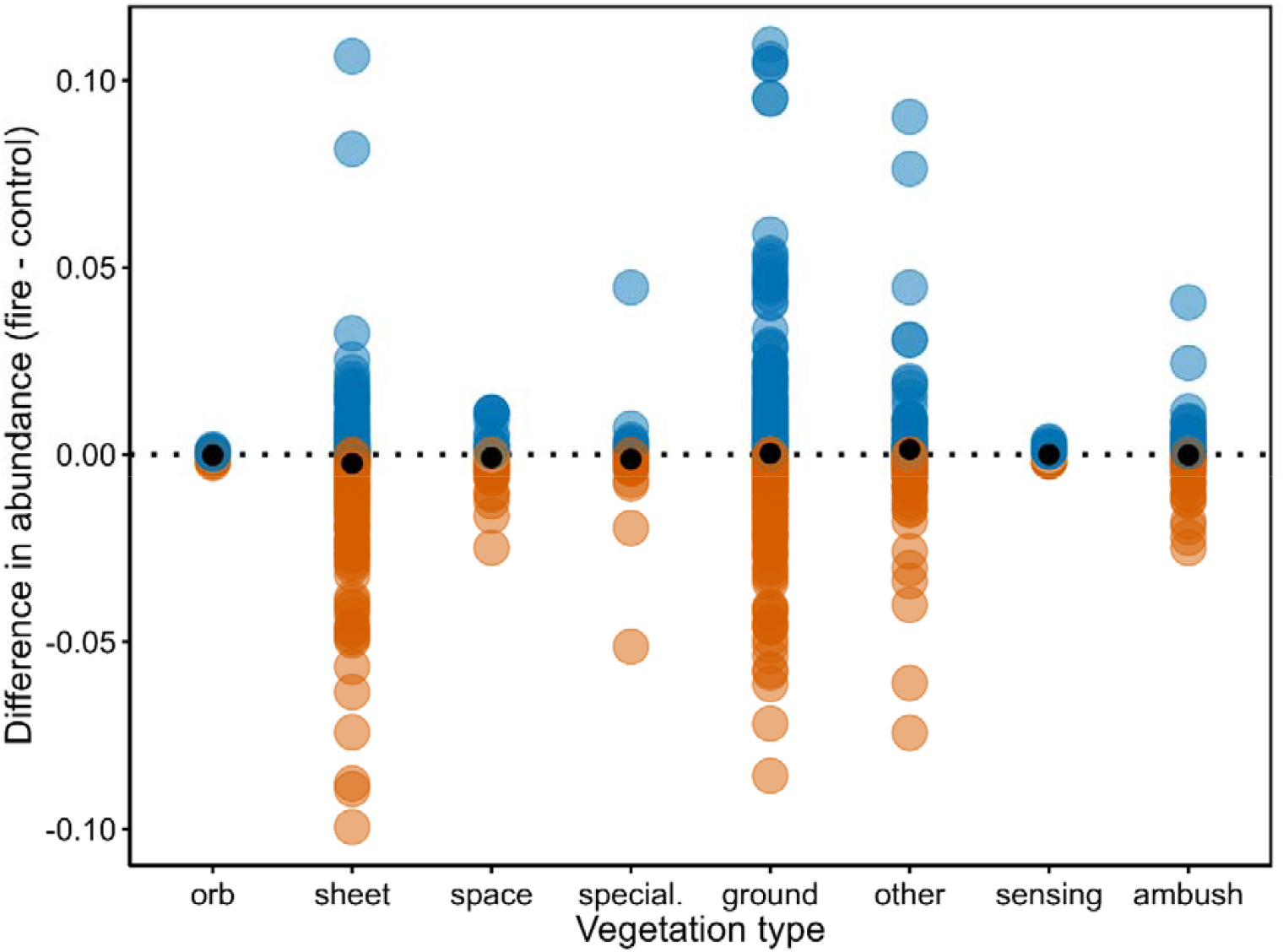
Difference in abundance between fire and control treatments for each hunting guild, for each species in each study. Blue = higher abundance in fire treatment, red = lower abundance in fire treatment, white = no difference, small black dot = mean difference in abundance for that guild. There was a significant difference between sheet web weavers and other hunters. Plot has been zoomed in to abundance differences between -0.1 and 0.1 for visualisation (N = 2595).

There was a significant relationship between presence/absence category and hunting guild (Model 3; marginal R^2^ = 0.021, conditional R^2^ = 0.214; Table 1). Both orb web weavers and sheet web weavers differed from ambush hunters, ground hunters, and other hunters in that a greater proportion of the two web weaving guilds “disappeared” and fewer “colonised” after fire (Figure 5; Table S7; Table S8). Time since fire and vegetation type also had a significant effect on presence/absence (Table 1; Table S7). Species that “disappeared” were recorded sooner after fire than those that “colonised” (Figure 6), while forests had more species that “disappeared” and “colonised” than grasslands (Figure 7).

**Figure 5.**
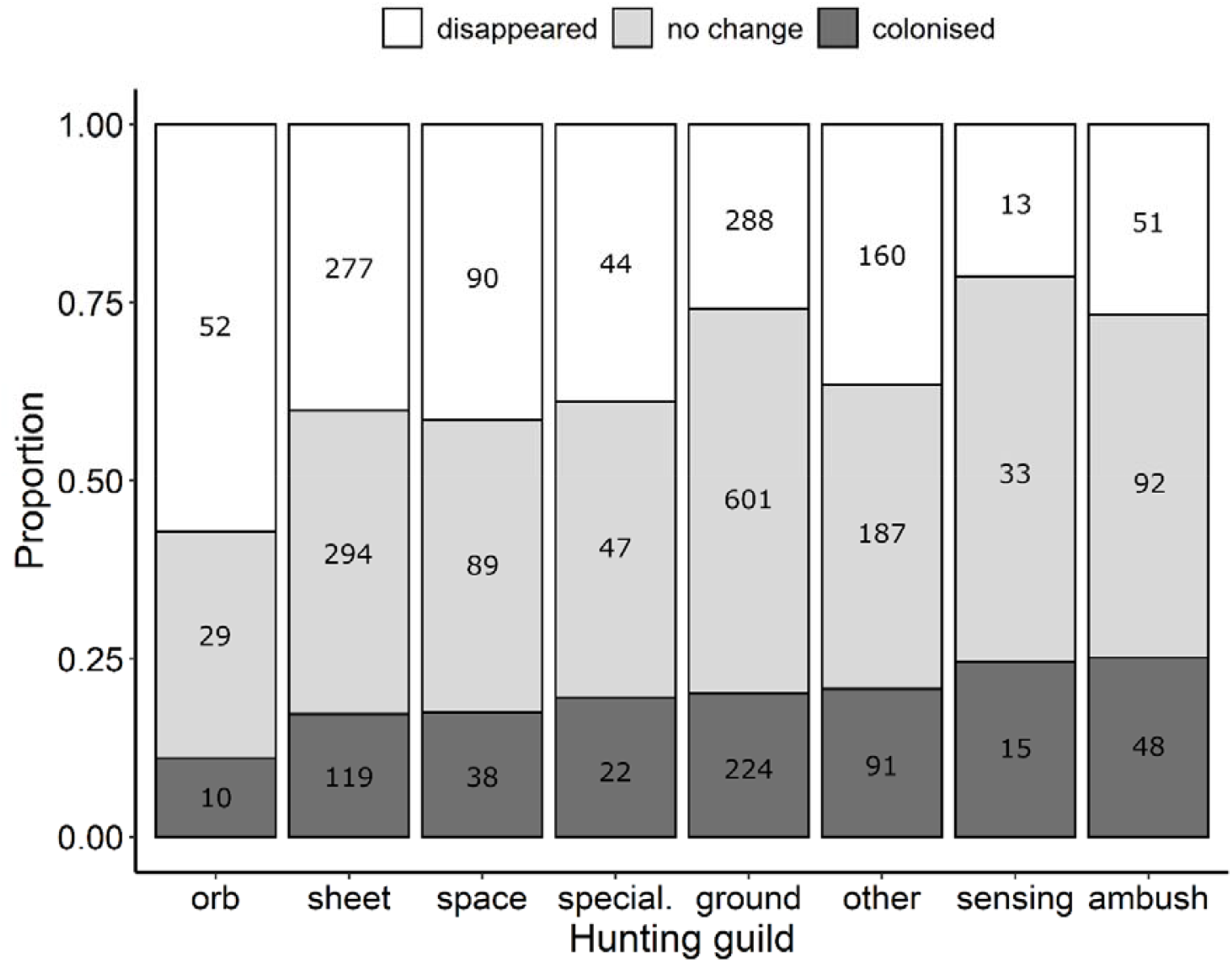
Proportion of datapoints (fire-control differences) for each hunting guild that fell into the three presence/absence categories: present in control treatment only (“disappeared”), present in both control and fire treatment (“no change”), present in fire treatment only (“colonised”). The total number of datapoints are shown within bars. There was a significant difference between both orb and sheet web weavers and ambush hunter, ground hunters, and other hunters (N = 2914).

**Figure 6.**
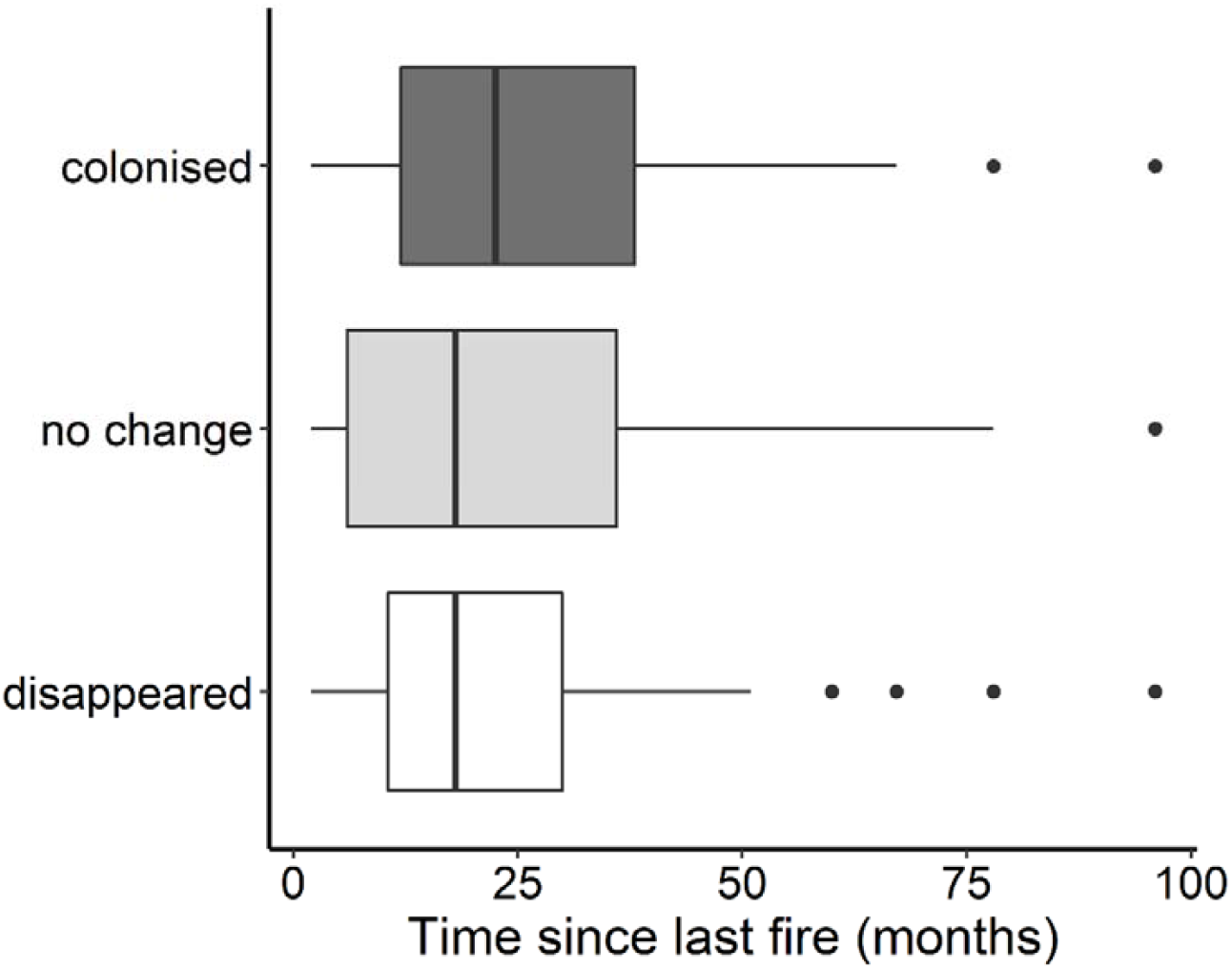
Relationship between time since last fire and the three presence/absence categories: present in control treatment only (“disappeared”), present in both control and fire treatment (“no change”), present in fire treatment only (“colonised”; N = 2914).

**Figure 7.**
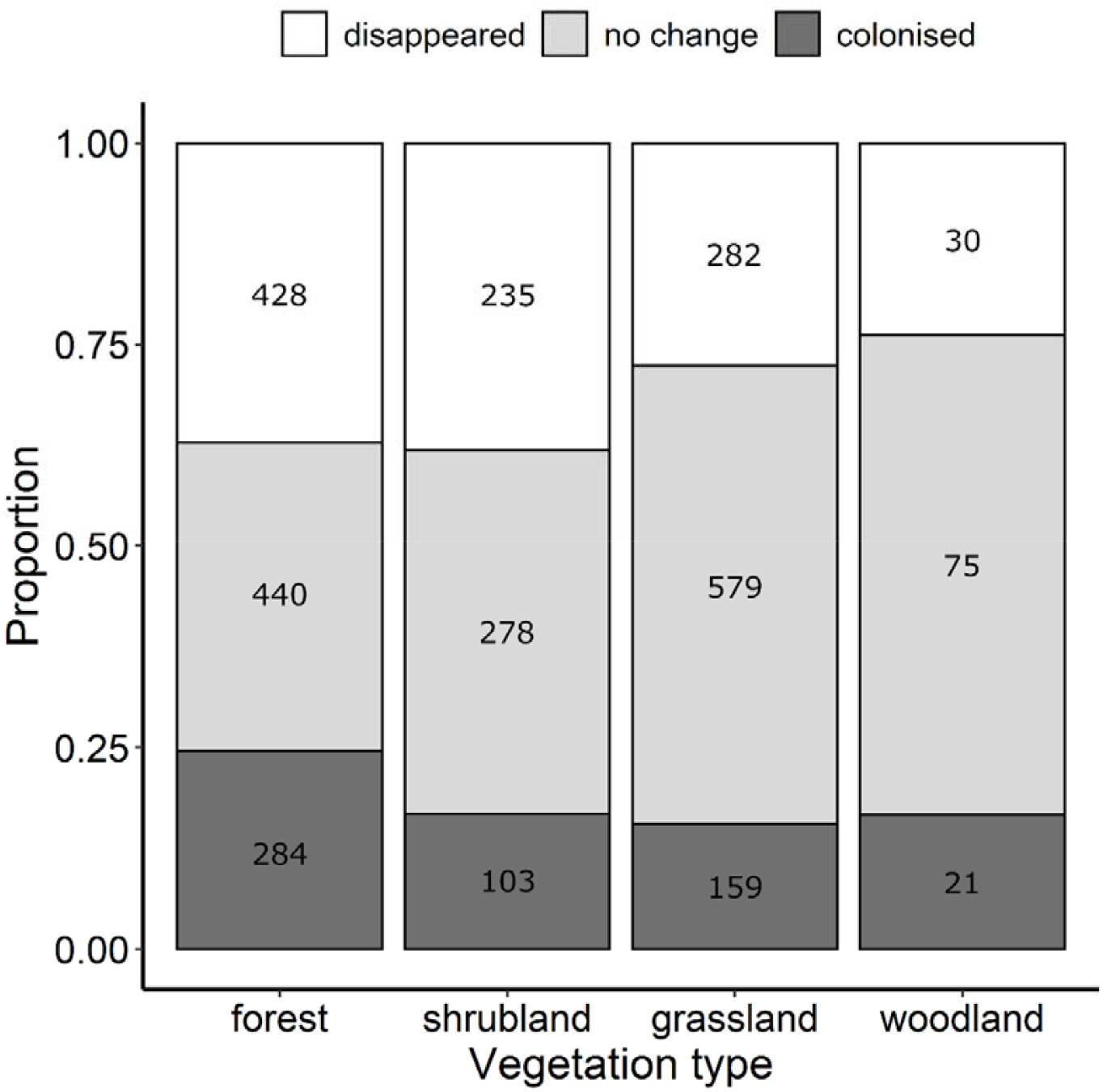
Proportion of datapoints (fire-control differences) for each vegetation type that fell into the three presence/absence categories: present in control treatment only (“disappeared”), present in both control and fire treatment (“no change”), present in fire treatment only (“colonised”). The total number of datapoints are shown within bars. There was a significant difference between forest and grassland vegetation types (N = 2914).

## DISCUSSION

We investigated the global impact of fire on ground-active spiders through a comparative analysis of published abundance and presence/absence data. Our results show that the effect of fire on spider families is independent of phylogenetic relatedness, and closely related families do not tend to show similar responses. In fact, we found that the effect of fire on individual families is highly variable among studies. It is therefore difficult to make broad generalisations about the impacts of fire on spiders at the family level; however, at the species level, we detected relationships between presence/absence categories and hunting guild, vegetation type, and time since fire. The relationship with time since fire highlights the importance of sampling across temporal scales. An understanding of this is essential to predict the impact of fire and increasing fire frequencies under climate change [5-7]. In particular, ecosystems may experience greater impacts when fire intervals are short, due to reduced time for re-establishment and for ecological succession to progress [61, 62].

It is important to note that our analyses reflect the available data, and studies testing the response of spiders to fire are limited. Additionally, the data are biased towards ground-dwelling taxa (which are captured most frequently in pitfall traps) but arboreal species contribute to species richness in many ecosystems and may be more influenced by changes to vegetation structure as a result of fire [63]. It was necessary to restrict our analysis to data obtained from pitfall trapping as it allowed us to standardise abundance within and between studies. However, studies that employ multiple sampling techniques designed to capture the full spider assemblage are likely to detect more informative patterns in spider responses to fire and life history traits underpinning those responses. We also acknowledge that pitfall trap design varied among studies (size of trap, preservation fluid or lack thereof, traps with or without a roof), which may influence the species recorded. Trap design was consistent among treatments within studies; however, it is possible that the efficacy of pitfall traps differed between unburned and burned habitat. For example, some species may be more attracted to pitfall traps with roofs in recently burned sites where vegetation cover is minimal. These potentially confounding factors should be considered when interpreting the results of this study.

Most spider families in our analysis showed both positive and negative responses to fire. This is likely due to differences in species assemblages and variation in other site-specific factors, such as species interactions, habitat structure, climate, and seasonality, as well as fire history variables that we were unable to include in our models (because they were not known/not reported), and other ecological disturbances (e.g., logging). Time since fire did have a significant effect on abundance for the family Lycosidae. Lycosids (wolf spiders) are ground-dwelling active hunters and burrowing habits and high dispersal ability (ballooning) have been recorded in some species [28]. Lycosids are often reported to increase in abundance following fire [39, 64, 65]. Our results showed a global pattern of higher abundance of lycosids in the fire treatment, but only from around two years after fire onwards. Prior to this, abundance was lower in the fire treatment. This suggests that fire may have direct impacts (mortality) but that burned habitats are beneficial to, or highly colonised by lycosids. Overall, the family level analysis detected some relationships with time since fire, but this taxonomic level is too coarse to investigate the predictors of response to fire.

Of the nearly 50,000 spider species currently recognised [26], trait data are available for fewer than 8000 species (16%) [28]. Given this, we only included hunting guild, ballooning, and female total body length in our analyses, three traits that are expected to influence response to fire. Our results suggest that a trait-based approach could increase predictive power when species level abundance or presence/absence data are analysed (as done in Models 2 and 3). We found that orb web weavers and sheet web weavers differed from ambush hunters, ground hunters, and other hunters in that a greater proportion of the two web weaving guilds “disappeared” and fewer “colonised” in the fire treatment. Fire also had a more negative effect on the abundance of sheet web weavers relative to other hunters. Web weaving spiders rely on vegetation for web anchor points [66, 67], and vegetation structure is affected by fire [68]. The higher proportion of species that “disappeared” may reflect a lack of suitable vegetation for web sites, or decreased habitat complexity in burned habitats [40, 41]. These guilds have been found to be associated with unburned sites in previous studies, for example, Malumbres-Olarte et al. [69] found that sheet web building spiders were associated with unburned sites in New Zealand grasslands. Conversely, ambush hunters often burrow or use retreat sites which may protect them during the fire event and/or be less altered in the post fire environment (soil and rock do not burn) [38], while ground and other hunters may benefit from increased hunting success in the more open habitat after fire [39]. Web attachment points are also critical for space web weavers. There was a possible trend of more species that “disappeared” in the space web weaver guild (Figure 5), but we did not detect significant differences to the other guilds. It is also important to note that orb web weavers are arboreal and therefore less likely to be captured in pitfall traps. However, this is unlikely to affect differences between unburned and burned sites within studies, unless orb web weavers move along the ground, and thus encounter pitfall traps, less in burned habitat, and it is unknown whether habitat changes post-fire affect spider behaviour. Overall, we detected a broad-scale relationship with hunting guild, but this is likely to interact with local site and fire specific factors.

Family and species level responses to fire also differed between vegetation types. Mean family level abundance was higher in the fire treatment for forests, but lower in the fire treatment for grassland, shrubland, and woodland habitats. Additionally, a higher proportion of species “disappeared” or “colonised” in the fire treatment for forest compared to grassland vegetation types, suggesting there may be a higher turnover of species in forests following fire. We did not analyse other diversity or community composition indices in this study, but this would likely give a more complete picture of the effects of fire. For example, de Brito et al. [70] found that arachnid abundance was lower, but diversity was higher in burned areas of Brazilian savannah. Similarly, Langlands et al. [71] found no difference in spider abundance across post-fire years, but significant changes in species composition in Australian spinifex habitat. Moretti et al. [72] found that spider community composition (but not overall abundance) varied between areas of Swiss chestnut forest experiencing different fire regimes. In all cases there were species unique to certain post-fire ages. Changes in abundance and/or species assemblages are predicted to be driven by changes to vegetation structure and habitat openness as the result of fire rather than fire *per se* [73, 74]. Therefore, the impacts of fire may be greater for assemblages from more complex habitats (i.e., forests) where habitat modification by fire is more pronounced. Additionally, fire behaviour differs between vegetation types, with grasslands and woodlands showing higher spread rates and intensities, more fine fuel biomass consumption, and increased release of heat when compared to forest ecosystems [75]. These differences in fire behaviour are likely to influence the extent of vegetation modification and have direct and indirect impacts on invertebrate taxa. The response of individual species to fire is also likely to be associated with finer details of vegetation structure that we were not able to test here.

Due to variation in study designs and limited reporting, we were not able to include other fire regime characteristics in our analysis, but these are likely to be key factors affecting how spiders respond to fire [42, 76]. For example, Andersen and Müller [77] found that spider abundance declined following late season, but not early season fires in northern Australia. Mason et al. [78] found that no mygalomorph spiders survived a high intensity wildfire, while nearly all survived a low intensity controlled burn in Western Australia. Additionally, Hanula and Wade [37] found that the effect of fire frequency differed among spider genera in pine stands in Florida. Many invertebrates also show seasonal variation in presence and abundance [79, 80]; therefore, the timing of sampling will have a major influence on species assemblages. Seasonality is difficult to incorporate into a global analysis due to geographic variation in seasons and the timing of fires; however, to minimise its impacts, we only included studies that surveyed control and fire treatments concurrently, or during the same season before and after fire. Where possible, studies should report detailed information about fire severity, intensity, frequency, season, size, and time since last fire, for both the treatment and control sites, and these should be included in analyses. Additionally, we recognise that replicated studies to test these factors are challenging to achieve when working with large-scale and often unpredictable disturbances such as fire, and suggest studies utilising future management burns (e.g., before-after-control-impact studies) may provide the most robust design [81, 82]. Given that time since fire was found to be an important factor, these studies should sample at multiple time points post-fire and consider each of those time points separately (i.e., not combine temporal sampling into one “after fire” group).

## CONCLUSIONS

More research is needed to understand the impacts of fire on spiders and invertebrates more broadly, and this will be most informative for management decisions when conducted at a local scale so that relationships with site-specific landscape and fire factors can be examined. As such, caution is needed when making broad-scale assumptions, but despite this we did find some global patterns in the response of spiders to fire and show that a trait-based approach may increase predictive power. Currently, this predictive ability is somewhat limited by a lack of knowledge, which highlights the importance of further taxonomic research and fundamental studies of the biology and ecology of invertebrate species. However, we did find evidence that hunting guilds differ in their response to fire, specifically that fire has a negative impact on orb and sheet web weavers. Additionally, short fire intervals may be a threat to some spiders, with abundance of lycosids initially lower after fire, and “disappearing” species detected sooner after fire than “colonising” species. This is important when considering the impact of changing fire regimes and designing fire management strategies to conserve biodiversity (e.g., determining fire intervals), and may help to identify priority species and ecosystems for more detailed study at a local scale. Finally, although not without limitations (noted above), we show that analyses of published raw data can be used to detect broad-scale ecological patterns and provide an alternative to traditional meta-analytical approaches, which is not limited by the questions asked in the original studies. We encourage authors to publish datasets along with their manuscripts so that their studies can be easily included in analyses such as this one.

## ACKNOWLEDGEMENTS

We acknowledge the authors of the original research analysed in this work.

## DATA ACCESSIBILITY

The datasets and code used during the current study are available from the Dryad Digital Repository: doi:xxxxxxx

## AUTHORS’ CONTRIBUTIONS

CAM, JM, and IM conceived of the research; CAM and RR performed the literature search; CAM extracted the abundance/presence data; CAM and JS compiled the trait data; CAM and IM designed and performed the comparative analysis; CAM and IM interpreted results; CAM wrote the manuscript, and all authors contributed to revisions.

