## Supplementary Information for "Assessing the impact of fire on spiders through a global comparative analysiss"

**This PDF file includes:**

Figures S1 to S6

Tables S1 to S8

Supplementary references

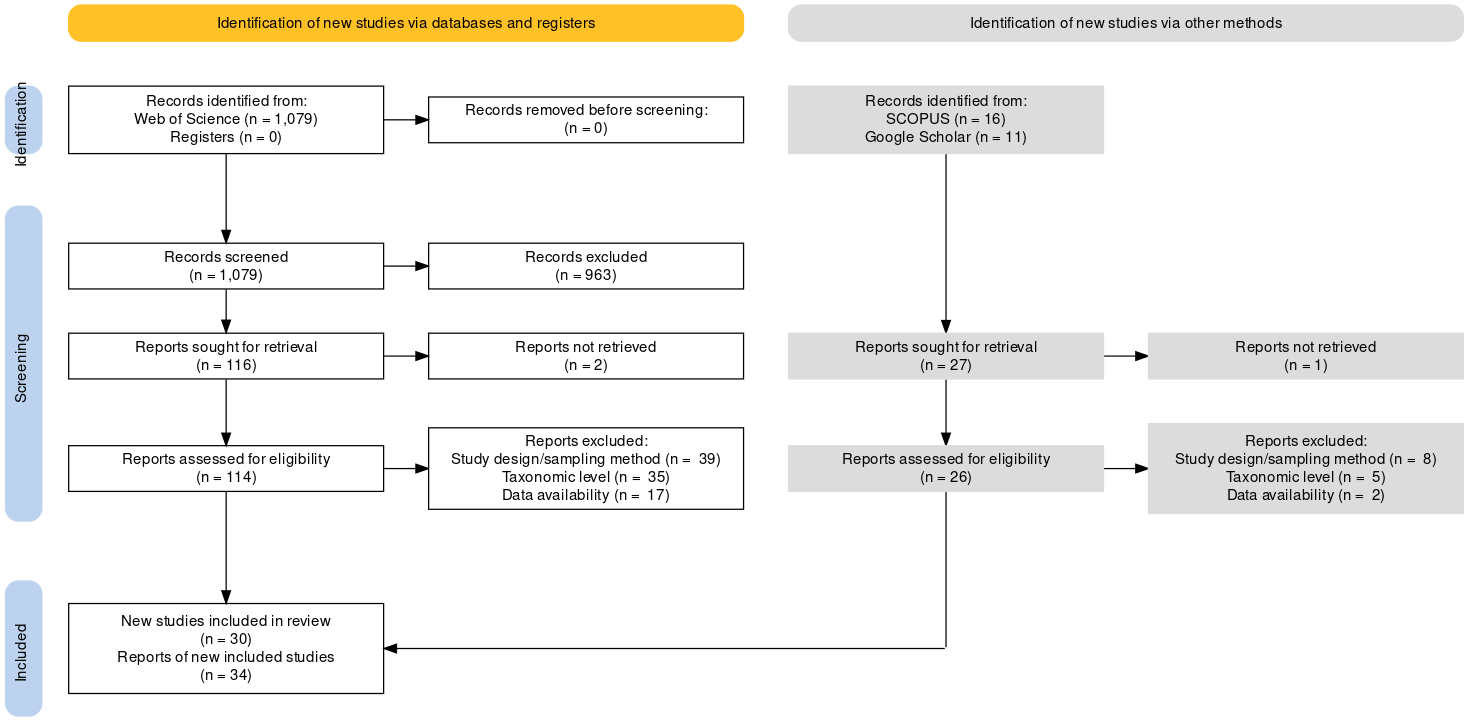

**Figure S1.** PRISMA 2020 (Haddaway *et al.* 2022) flow diagram of the literature search employed for this study.

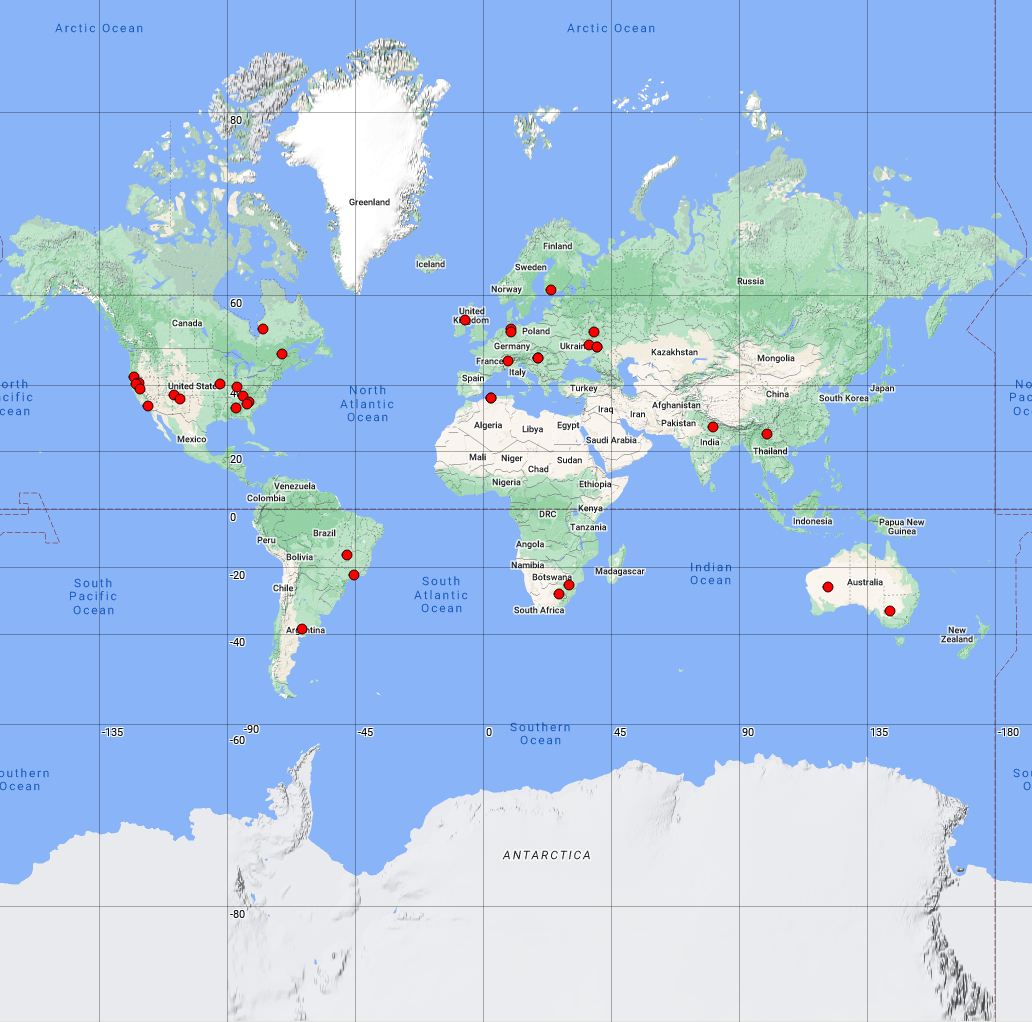

**Figure S2.** Map showing locations of the 34 studies (30 datasets) included in our analysis. See Table S1 for further details including GPS coordinates. Map created using Map Maker ([https://maps.co](https://hes32-ctp.trendmicro.com/wis/clicktime/v1/query?url=https%3a%2f%2fmaps.co%2fml%2fl%2fEiWwa0cMfnLsUZ94FSNeDQ%2fWGV4jvgBcCk20yqjK5jUKw%2fbMAoLMG10WfO8nv5Kg9HnA&umid=841add70-4664-40fd-b77d-2f13b32460f0&auth=89a422ce48cf9afc268cabe806cc53ea452e36bd-9e799e48e3723d38de1b74420b1e3a3408cf5420)).

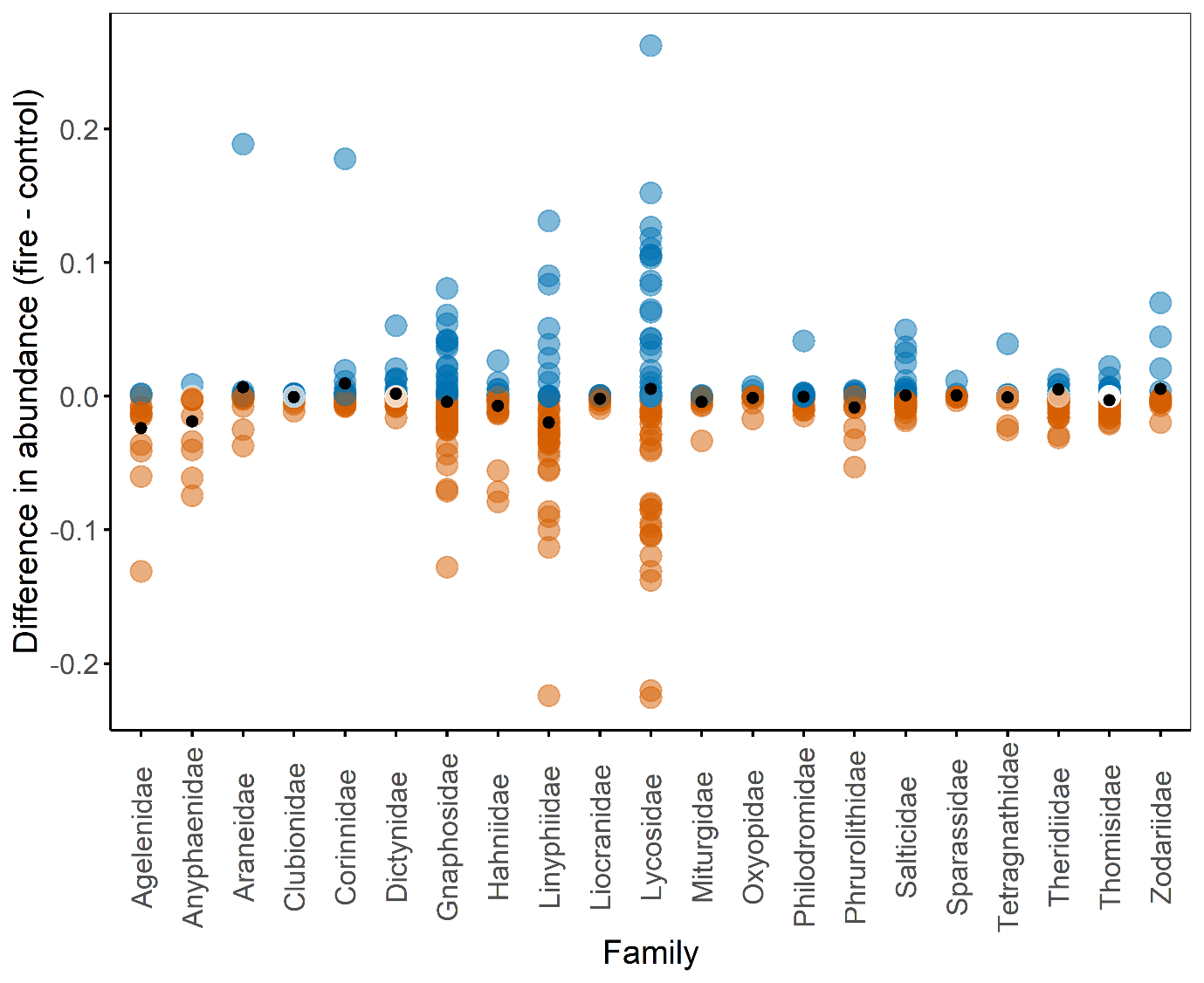

**Figure S3.** Difference in abundance between fire and control treatments for each family, for each study. Blue = higher abundance in fire treatment, red = lower abundance in fire treatment, white = no difference, small black dot = mean difference in abundance for that family. Only data for families with ≥10 datapoints are presented. Four outlying datapoints were removed for visualisation (N = 564).

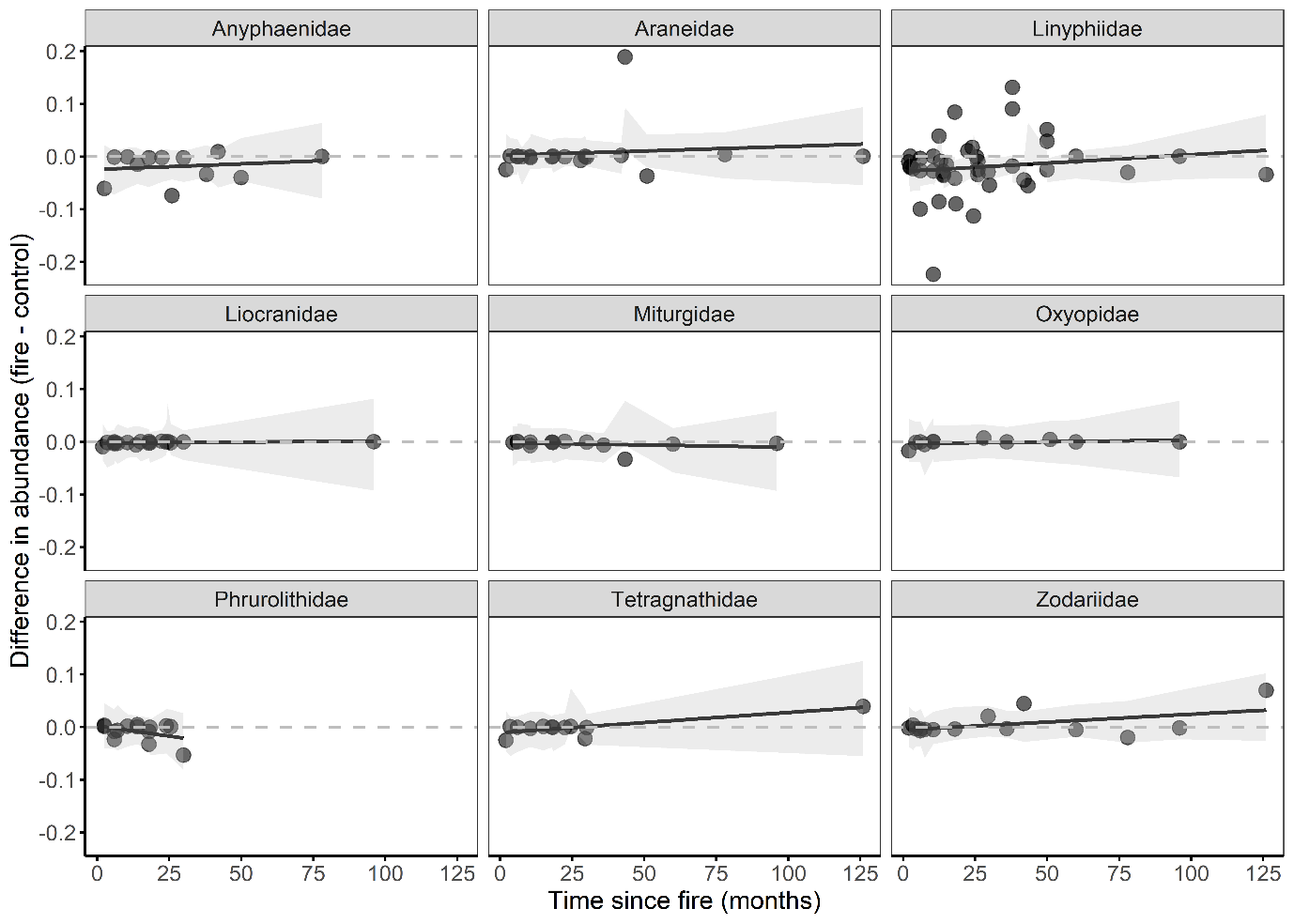

**Figure S4.** Relationship between time since fire (months) and difference in abundance between fire and control treatment within each family (Model 1). This figure shows the families not included in Figure 3. Grey shading = 95% confidence bounds (N = 564).

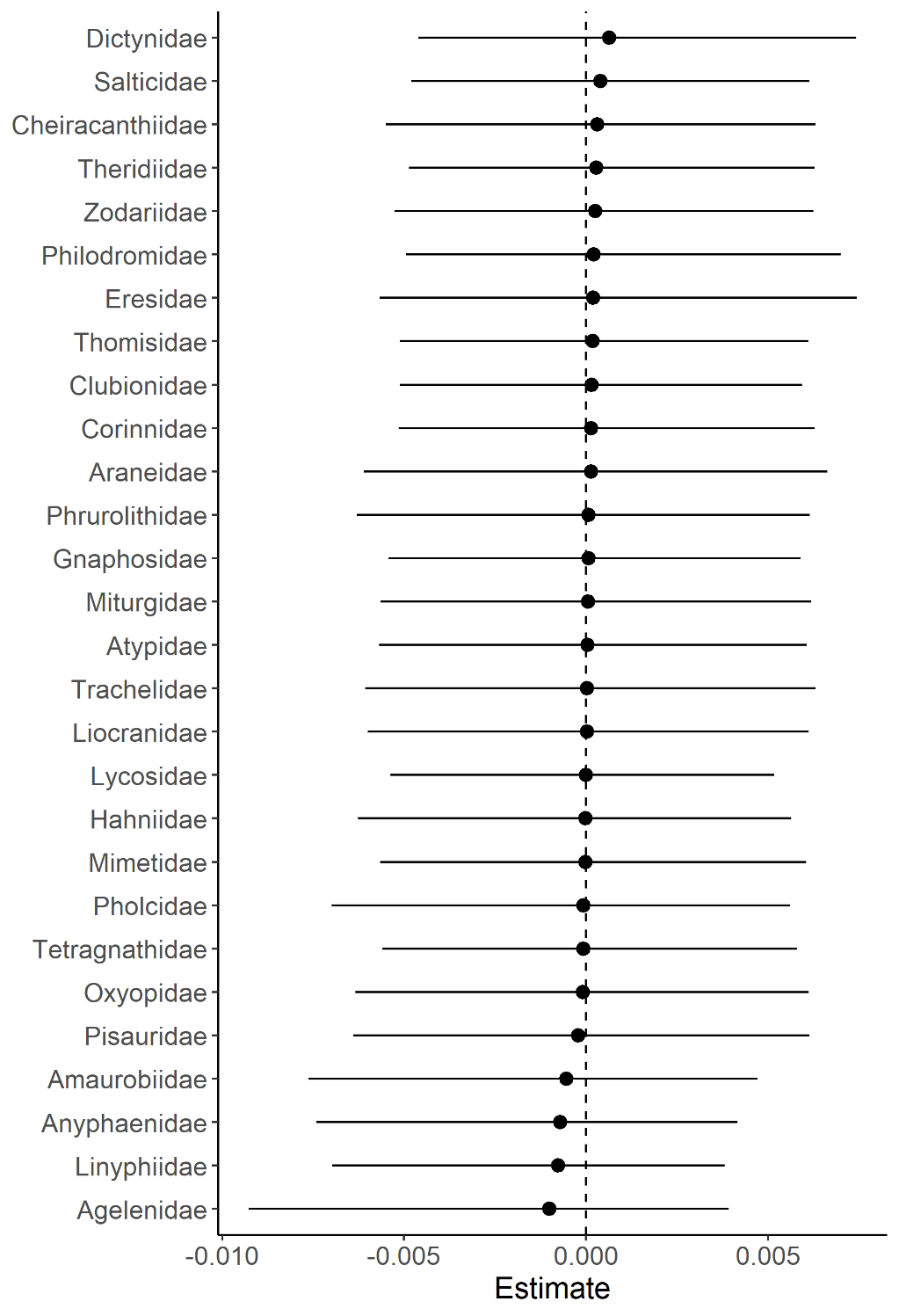

**Figure S5.** Random effect of family from the intercept only model (difference in abundance ~ 1) with phylogeny, family, and study ID as random factors. Only families with ≥10 datapoints are included (N = 564).

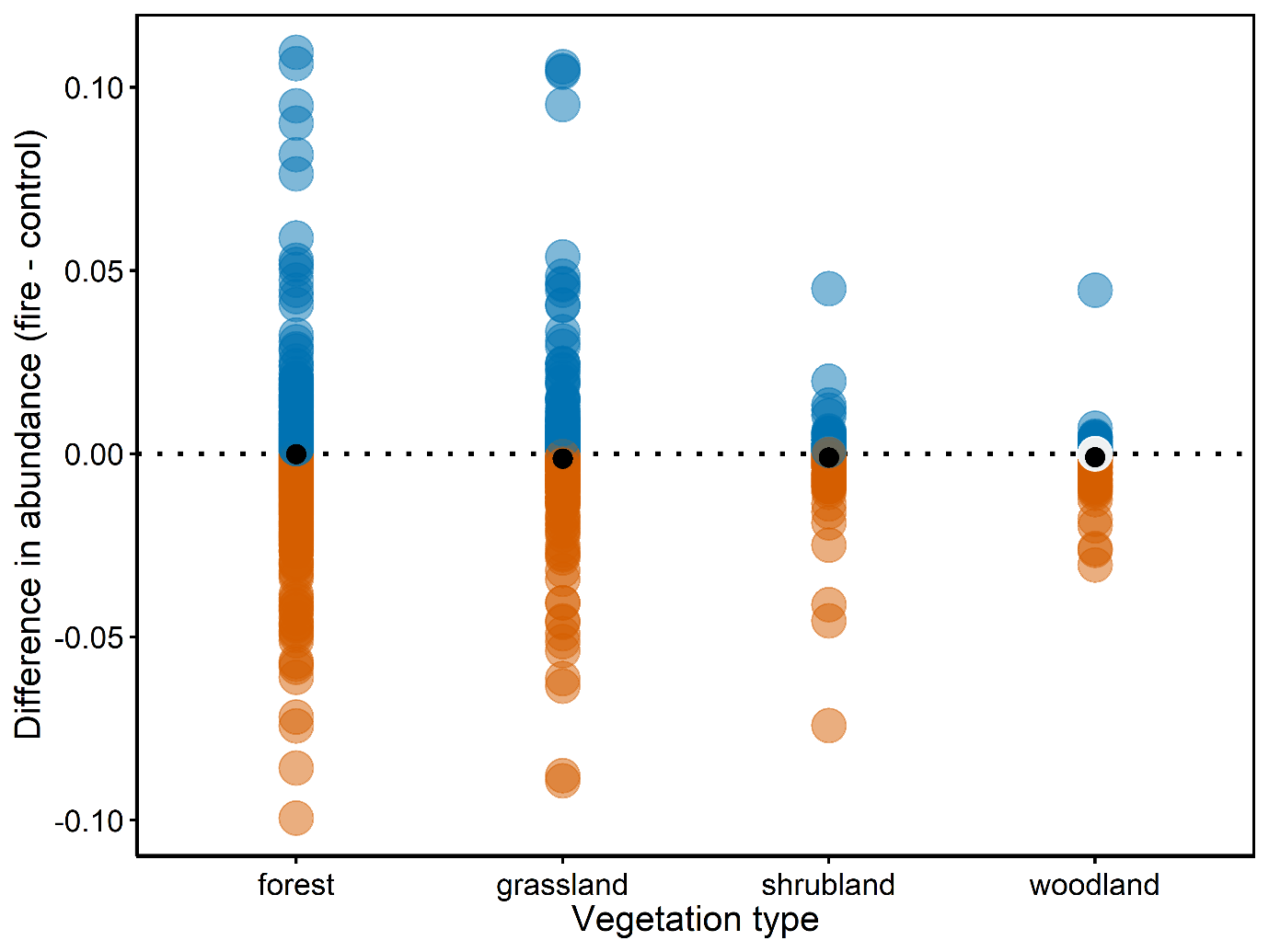
**Figure S6.** Difference in abundance between fire and control treatments for each vegetation type, for each species in each study. Blue = higher abundance in fire treatment, red = lower abundance in fire treatment, white = no difference, small black dot = mean difference in abundance for that vegetation type. Plot has been zoomed in to abundance differences between -0.1 and 0.1 for visualisation (N = 2595).

**Table S1.** Summary of the 34 studies (30 datasets) included in our analysis. Design: UB = unburned vs burned, BA = before vs after; Sample effort: (plots*traps*days for the control treatment + plot*traps*days for the fire treatment)/2; Notes on data use: description of the data used for our analysis; Ab: abundance data were available for this study; P/A: presence/absence data were available for this study.

| **Study ID** | **Location** | **Design** | **Veg. type** | **Latitude** | **Longitude** | **Sample effort** | **Notes on data use** | **Ab.** | **P/A** | **Reference** |
| --- | --- | --- | --- | --- | --- | --- | --- | --- | --- | --- |
| C | Brazil | UB | forest | -22.6586 | -45.4428 | 90 | Used reforested Araucaria plots. | × | × | Baretta *et al.* (2007) |
| I | Brazil | UB | woodland | -15.9392 | -47.8853 | 2280 | Used quarterly burned. For other treatments, most sampling was pre-fire. | × | × | De Brito Freire-Jr and Motta (2011) |
| L | U.S.A. | UB | forest | 40.50017 | -121.001 | 940 | Used unlogged sites. | × | × | Gillette *et al.* (2008) |
| M | U.S.A. | UB | forest | 35.28583 | -82.3283 | 1176 | Family level data only. | × |  | Greenberg *et al.* (2010) |
| N | South Africa | UB | grassland | -28.4914 | 26.8088 | 10920 |  | × | × | Haddad *et al.* (2015) |
| Q | U.S.A. | UB | grassland | 40.17361 | -92.5833 | 672 |  | × | × | Haskins and Shaddy (1986) |
| R | U.S.A. | UB | woodland | 37.23095 | -108.463 | 9450 | Used Chapin Mesa. | × | × | Higgins *et al.* (2014) |
| S | India | UB | grassland | 27.81667 | 81.01667 | 5010 |  | × | × | Hore and Uniyal (2008) |
| V | South Africa | UB | grassland | -25.935 | 30.37722 | 972 | Used communal land, annual farmland as they had similar grazing intensity. | × | × | Jansen *et al.* (2013) |
| W | Finland | UB | forest | 60.73333 | 23.75 | 1410 |  | × | × | Koponen (2003) |
| X | *as above* | “ | “ | “ | “ | “ |  | × | × | Koponen (2005) |
| Y | Germany | UB | shrubland | 53.25 | 9.966667 | 6864 |  | × | × | Krause and Assmann (2016) |
| Z | Australia | UB | woodland | -33.7167 | 143.0333 | 1120 | Family level data only. | × |  | Kwok and Eldridge (2015) |
| AB | Australia | UB | shrubland | -26.35 | 121.3333 | 7710 |  | × | × | Langlands *et al.* (2011) |
| AC | *as above* | “ | “ | “ | “ | “ | Family level data only. | × |  | Langlands *et al.* (2012) |
| AD | Canada | UB | forest | 47.68333 | -70.6833 | 738 | Used undisturbed sites. | × | × | Larrivée *et al.* (2005) |
| AF | China | UB | forest | 25.56667 | 99.91667 | 9100 |  | × | × | Ma *et al.* (2014) |
| AG | *as above* | “ | “ | “ | “ | “ | Family level data only. Used unburned, 2- and 10-year sites. | × |  | Ma *et al.* (2013) |
| AK | Algeria | UB | forest | 36.42944 | 2.91388 | 7300 |  |  | × | Mansouri *et al.* (2019) |
| AL | Ireland | UB | shrubland | 55.18086 | -6.23642 | 4932 | Used ungrazed sites. | × | × | McFerran *et al.* (1995) |
| AN | Switzerland | UB | forest | 46.15 | 8.7333 | 2556 |  | × | × | Moretti (2000) |
| AO | *as above* | “ | “ | “ | “ | “ | Used recent fire, single fire 1-2 years, repeated fire 2-3 years. | × | × | Moretti *et al.* (2002) |
| AS | Ukraine | UB | grassland | 49.28333 | 40.0 | 3600 | Used 1 and 3 years after fire. |  | × | Polchaninova (2015) |
| AT | Ukraine | BA | grassland | 49.9304 | 37.30956 | 430 | Used before and after, then burned and unburned for subsequent years. All data from Autumn. | × | × | Polchaninova *et al.* (2019) |
| AU | Russia | UB | grassland | 52.59773 | 38.92434 | 660 | Used feather grass forbe steepe control. | × | × | Polchaninova *et al.* (2016) |
| AV | Argentina | UB | woodland | -38.7667 | -63.75 | 560 |  | × | × | Pompozzi *et al.* (2011) |
| AZ | Hungary | UB | grassland | 46.78333 | 19.45 | 11240 |  |  | × | Samu *et al.* (2010) |
| BA | Germany | UB | forest | 52.70008 | 10.00331 | 2192 | Used 1976 and 1977. | × | × | Schaefer (1980) |
| BB | U.S.A. | BA | shrubland | 34.10919 | -117.712 | 360 | Used Spring before and after fire. | × | × | Spear *et al.* (2017) |
| BC | U.S.A. | UB | woodland | 40.10285 | -122.047 | 504 | Family level data only. | × |  | Underwood and Quinn (2010) |
| BG | U.S.A. | UB | forest | 34.73195 | -82.8518 | 720 | Family level data only. | × |  | Vickers and Culin (2014) |
| BI | Canada | UB | forest | 53.38333 | -77.5 | 540 | Family level data only. | × |  | Koponen (1993) |
| BJ | U.S.A. | UB | forest | 35.92 | -106.521 | 2214 |  | × | × | Brantley (2020) |
| BK | U.S.A | BA | forest | 39.32579 | -86.4199 | 3360 |  | × | × | Milne *et al.* (2021) |

**Table S2.** Inverse Wishart priors used for each of the phylogenetic mixed models. We changed the random effects nu value for Model 1 due to convergence issues. Prior choice did not affect results.

| **Model** | **Residual (R)** | | | **Random Effects (G)** | | | |
| --- | --- | --- | --- | --- | --- | --- | --- |
|  | **V** | **nu** | **fix** | **V** | **nu** | **alpha.mu** | **alpha.V** |
| 1 | 1 | 0.002 |  | 1 | 2 | 0 | 25^2 |
| 2 | 1 | 0.002 |  | 1 | 0.002 | 0 | 25^2 |
| 3 | 1 |  | 1 | 1 | 0.002 | 0 | 25^2 |

**Table S3.** Association between the family level difference in abundance between fire and control treatments and family, vegetation type (vegetation), fire type, absolute latitude (latitude), time since fire (months) and the interaction between family and time since fire (family × months; Model 1; N = 564).

|  | **Difference in abundance** | | |
| --- | --- | --- | --- |
| **Predictors** | **Lower** | **Upper** | **P** |
| Family (Agelenidae) | -11.6 | 9.65 | 0.989 |
| Family (Anyphaenidae) | -8.23 | 11.8 | 0.976 |
| Family (Araneidae) | -12.1 | 13.0 | 0.952 |
| Family (Clubionidae) | -10.2 | 9.71 | 0.981 |
| Family (Corinnidae) | -10.8 | 8.79 | 0.998 |
| Family (Dictynidae) | -9.22 | 11.4 | 0.988 |
| Family (Gnaphosidae) | -10.5 | 9.78 | 0.991 |
| Family (Hahniidae) | -10.5 | 10.2 | 0.984 |
| Family (Linyphiidae) | -12.1 | 12.4 | 0.977 |
| Family (Liocranidae) | -9.16 | 10.4 | 0.984 |
| Family (Miturgidae) | -10.6 | 9.16 | 0.996 |
| Family (Oxyopidae) | -8.78 | 9.15 | 0.997 |
| Family (Philodromidae) | -10.3 | 9.93 | 0.982 |
| Family (Phrurolithidae) | -9.67 | 9.93 | 0.991 |
| Family (Salticidae) | -9.98 | 10.3 | 0.976 |
| Family (Sparassidae) | -9.89 | 10.5 | 0.996 |
| Family (Tetragnathidae) | -12.2 | 11.6 | 0.963 |
| Family (Theridiidae) | -12.7 | 12.4 | 0.982 |
| Family (Thomisidae) | -8.70 | 7.75 | 0.984 |
| Family (Zodariidae) | -0.113 | 9.70 | 0.989 |
| Months | **2.86E^-04^** | **1.16E^-03^** | **0.002** |
| Vegetation (grassland) | -0.023 | 1.30E^-03^ | 0.066 |
| Vegetation (shrubland) | -0.031 | 7.26E^-03^ | 0.212 |
| Vegetation (woodland) | **-0.038** | **7.99E^-04^** | **0.046** |
| Fire type | -0.013 | 0.016 | 0.980 |
| Latitude | -3.98E^-04^ | 8.68E^-04^ | 0.382 |
| Family (Agelenidae) × months | **-1.97E^-03^** | **-4.58E^-04^** | **0.003** |
| Family (Anyphaenidae) × months | -1.89E^-03^ | 8.27E^-04^ | 0.390 |
| Family (Araneidae) × months | -1.43E^-03^ | 5.99E^-05^ | 0.062 |
| Family (Clubionidae) × months | **-1.42E^-03^** | **-8.35E^-05^** | **0.032** |
| Family (Corinnidae) × months | -1.31E^-03^ | 3.33E^-05^ | 0.045 |
| Family (Dictynidae) × months | **-1.35E^-03^** | **-6.52E^-05^** | **0.042** |
| Family (Gnaphosidae) × months | **-1.29E^-03^** | **-8.68E^-05^** | **0.028** |
| Family (Hahniidae) × months | **-1.61E^-03^** | **-2.48E^-04^** | **0.013** |
| Family (Linyphiidae) × months | -9.74E^-04^ | 2.63E^-04^ | 0.246 |
| Family (Liocranidae) × months | -1.64E^-03^ | 3.04E^-04^ | 0.179 |
| Family (Miturgidae) × months | -1.68E^-03^ | 2.01E^-04^ | 0.115 |
| Family (Oxyopidae) × months | -1.60E^-03^ | 1.63E^-04^ | 0.131 |
| Family (Philodromidae) × months | **-1.46E^-03^** | **-6.17E^-05^** | **0.031** |
| Family (Phrurolithidae) × months | -4.18E^-03^ | 1.11E^-03^ | 0.218 |
| Family (Salticidae) × months | **-1.30E^-03^** | **-2.04E^-05^** | **0.054** |
| Family (Sparassidae) × months | **-1.47E^-03^** | **-8.96E^-06^** | **0.057** |
| Family (Tetragnathidae) × months | -1.18E^-03^ | 4.21E^-04^ | 0.358 |
| Family (Theridiidae) × months | **-1.31E^-03^** | **-4.85E^-05^** | **0.039** |
| Family (Thomisidae) × months | **-1.34E^-03^** | **-1.27E^-04^** | **0.018** |
| Family (Zodariidae) × months | -1.11E^-03^ | 2.73E^-04^ | 0.231 |

Lower and upper represent the 95% confidence bounds from the posterior distribution of the estimate. Cases where the 95% confidence interval does not overlap zero, and therefore provides evidence of a significant effect, are highlighted in bold. Family (Lycosidae), and Vegetation (forest) were used for comparison.

**Table S4.** Comparison of family-level intercept only models with different random effect structures (N = 564).

| **Random Factors** | **df** | **DIC** | **ΔDIC** | **DIC weight** |
| --- | --- | --- | --- | --- |
| Study ID | 4 | -2026.3 | 0.00 | 0.562 |
| Family, Study ID | 5 | -2025.0 | 1.30 | 0.294 |
| Phylogeny, Family, Study ID | 6 | -2023.6 | 2.71 | 0.145 |

**Table S5.** Association between the species level difference in abundance between fire and control treatments and hunting guild (guild), ballooning, female body length (body length), vegetation type (vegetation), fire type, absolute latitude (latitude) and time since fire (months; Model 2; N = 2595). The complete case analysis was performed to show that data imputation did not affect inferences about female body length (imputed variable; N = 1812).

|  |  | **Difference in abundance** | | |
| --- | --- | --- | --- | --- |
| **Dataset** | **Predictors** | **Lower** | **Upper** | **P** |
| Imputed | Guild (ambush hunters) | 7.19E^-06^ | 8.08E^-03^ | 0.059 |
|  | Guild (ground hunters) | -2.26E^-04^ | 5.58E^-03^ | 0.059 |
|  | Guild (orb web weavers) | -1.09E^-03^ | 7.60E^-03^ | 0.116 |
|  | Guild (other hunters) | **1.12E^-03^** | **7.13E^-03^** | **0.006** |
|  | Guild (sensing web weavers) | -1.86E^-03^ | 9.50E^-03^ | 0.181 |
|  | Guild (space web weavers) | -8.84E^-04^ | 5.22E^-03^ | 0.161 |
|  | Guild (specialists) | -2.78E^-03^ | 5.40E^-03^ | 0.513 |
|  | Ballooning | -2.73E^-03^ | 2.11E^-03^ | 0.871 |
|  | Body length | -3.31E^-04^ | 1.08E^-04^ | 0.265 |
|  | Vegetation (grassland) | -4.73E^-03^ | 9.63E^-05^ | 0.086 |
|  | Vegetation (shrubland) | -4.90E^-03^ | 1.49E^-03^ | 0.270 |
|  | Vegetation (woodland) | -7.07E^-03^ | 1.51E^-03^ | 0.201 |
|  | Fire type | -3.53E^-03^ | 2.07E^-03^ | 0.597 |
|  | Latitude | -1.39E^-04^ | 9.85E^-05^ | 0.619 |
|  | Months | -1.00E^-05^ | 4.74E^-05^ | 0.197 |
| Complete case | Guild (ambush hunters) | -4.82E^-04^ | 1.02E^-02^ | 0.096 |
|  | Guild (ground hunters) | -8.23E^-04^ | 7.44E^-03^ | 0.116 |
|  | Guild (orb web weavers) | -1.59E^-03^ | 1.07E^-02^ | 0.178 |
|  | Guild (other hunters) | **4.95E^-04^** | **8.67E^-03^** | **0.018** |
|  | Guild (sensing web weavers) | -5.94E^-03^ | 1.28E^-02^ | 0.441 |
|  | Guild (space web weavers) | -2.61E^-03^ | 7.22E^-03^ | 0.350 |
|  | Guild (specialists) | -7.40E^-03^ | 6.26E^-03^ | 0.787 |
|  | Ballooning | -5.75E^-03^ | 3.22E^-03^ | 0.535 |
|  | Body length | -3.12E^-04^ | 1.99E^-04^ | 0.519 |
|  | Vegetation (grassland) | **-5.83E^-03^** | **-4.44E^-04^** | **0.009** |
|  | Vegetation (shrubland) | -6.23E^-03^ | 2.16E^-03^ | 0.289 |
|  | Vegetation (woodland) | -9.43E^-03^ | 3.43E^-03^ | 0.443 |
|  | Fire type | -5.03E^-03^ | 2.25E^-03^ | 0.508 |
|  | Latitude | -1.87E^-04^ | 1.35E^-04^ | 0.724 |
|  | Months | -5.69E^-07^ | 9.35E^-05^ | 0.049 |

Lower and upper represent the 95% confidence bounds from the posterior distribution of the estimate. Cases where the 95% confidence interval does not overlap zero, and therefore provides evidence of a significant effect, are highlighted in bold. Guild (sheet web weavers), and Vegetation (forest) were used for comparison.

**Table S6.** *Post hoc* pairwise comparisons between each of the hunting guilds based on the imputed dataset model in Table S5.

| **Response** | **Comparison** | **Lower** | **Upper** |
| --- | --- | --- | --- |
| Difference in abundance (fire - control) | Orb – Ambush | -5.69E-03 | 4.71E-03 |
|  | Orb – Ground | -3.84E-03 | 4.95E-03 |
|  | Orb – Other | -4.45E-03 | 4.82E-03 |
|  | Orb – Sensing | -6.51E-03 | 6.42E-03 |
|  | Orb – Sheet | -9.72E-04 | 8.21E-03 |
|  | Orb – Space | -3.56E-03 | 6.12E-03 |
|  | Orb – Specialist | -2.99E-03 | 7.60E-03 |
|  | Ambush – Ground | -2.49E-03 | 4.04E-03 |
|  | Ambush – Other | -3.63E-03 | 3.22E-03 |
|  | Ambush – Sensing | -6.60E-03 | 5.19E-03 |
|  | Ambush – Sheet | -3.42E-04 | 7.51E-03 |
|  | Ambush – Space | -2.63E-03 | 5.62E-03 |
|  | Ambush – Specialist | -2.24E-03 | 7.12E-03 |
|  | Ground – Other | -3.34E-03 | 1.03E-03 |
|  | Ground – Sensing | -6.75E-03 | 3.66E-03 |
|  | Ground – Sheet | -4.76E-04 | 5.71E-03 |
|  | Ground – Space | -2.57E-03 | 3.72E-03 |
|  | Ground – Specialist | -2.16E-03 | 4.56E-03 |
|  | Other – Sensing | -5.70E-03 | 5.53E-03 |
|  | Other – Sheet | **9.56E-04** | **6.94E-03** |
|  | Other – Space | -1.38E-03 | 4.97E-03 |
|  | Other – Specialist | -1.59E-03 | 5.99E-03 |
|  | Sensing – Sheet | -2.23E-03 | 9.30E-03 |
|  | Sensing – Space | -4.36E-03 | 7.46E-03 |
|  | Sensing – Specialist | -2.77E-03 | 8.71E-03 |
|  | Sheet – Space | -5.45E-03 | 8.34E-04 |
|  | Sheet – Specialist | -5.62E-03 | 2.65E-03 |
|  | Space – Specialist | -3.42E-03 | 5.21E-03 |

Cases where the 95% confidence interval does not overlap zero, and therefore provides evidence of a significant effect, are highlighted in bold

**Table S7.** Association between species presence/absence categories (“disappeared”, “no change”, “colonised”) and hunting guild (guild), ballooning, female body length (body length), vegetation type (vegetation), fire type, absolute latitude (latitude) and time since fire (months; Model 3; N = 2914). The complete case analysis was performed to show that data imputation did not affect inferences about female body length (imputed variable; N = 2108).

|  |  | **Difference in presence/absence** | | |
| --- | --- | --- | --- | --- |
| **Dataset** | **Predictors** | **Lower** | **Upper** | **P** |
| Imputed | Guild (ambush hunters) | **0.254** | **2.295** | **0.021** |
|  | Guild (ground hunters) | **0.316** | **1.999** | **0.003** |
|  | Guild (other hunters) | **0.317** | **1.917** | **0.003** |
|  | Guild (sensing web weavers) | -0.256 | 1.871 | 0.127 |
|  | Guild (sheet web weavers) | -0.294 | 1.318 | 0.206 |
|  | Guild (space web weavers) | -0.136 | 1.699 | 0.071 |
|  | Guild (specialists) | -0.350 | 1.481 | 0.258 |
|  | Ballooning | -0.697 | 0.101 | 0.141 |
|  | Body length | -0.034 | 0.021 | 0.593 |
|  | Vegetation (grassland) | **-0.461** | **-0.058** | **0.012** |
|  | Vegetation (shrubland) | -0.686 | 0.288 | 0.361 |
|  | Vegetation (woodland) | -1.064 | 0.108 | 0.104 |
|  | Fire type | -0.390 | 0.376 | 0.886 |
|  | Latitude | -0.023 | 0.010 | 0.441 |
|  | Months | **0.004** | **0.010** | **0.0007** |
| Complete case | Guild (ambush hunters) | **0.132** | **2.530** | **0.031** |
|  | Guild (ground hunters) | **0.236** | **2.108** | **0.007** |
|  | Guild (other hunters) | **0.002** | **1.826** | **0.034** |
|  | Guild (sensing web weavers) | -1.136 | 1.267 | 0.778 |
|  | Guild (sheet web weavers) | -0.554 | 1.162 | 0.493 |
|  | Guild (space web weavers) | -0.287 | 1.746 | 0.153 |
|  | Guild (specialists) | -0.787 | 1.330 | 0.690 |
|  | Ballooning | -0.814 | 0.277 | 0.267 |
|  | Body length | -0.022 | 0.034 | 0.679 |
|  | Vegetation (grassland) | **-0.417** | **-0.006** | **0.031** |
|  | Vegetation (shrubland) | -0.463 | 0.473 | 0.905 |
|  | Vegetation (woodland) | -0.979 | 0.401 | 0.382 |
|  | Fire type | -0.253 | 0.541 | 0.507 |
|  | Latitude | -0.027 | 0.012 | 0.539 |
|  | Months | **0.006** | **0.014** | **0.0007** |

Lower and upper represent the 95% confidence bounds from the posterior distribution of the estimate. Cases where the 95% confidence interval does not overlap zero, and therefore provides evidence of a significant effect, are highlighted in bold. Guild (orb web weavers), and Vegetation (forest) were used for comparison.

**Table S8.** *Post hoc* pairwise comparisons between each of the hunting guilds based on the imputed dataset model in Table SX.

| **Response** | **Comparison** | **Lower** | **Upper** |
| --- | --- | --- | --- |
| Difference in presence/absence  (3 categories: disappeared, no change, colonised) | Orb – Ambush | **-2.295** | **-0.254** |
|  | Orb – Ground | **-1.999** | **-0.316** |
|  | Orb – Other | **-1.917** | **-0.317** |
|  | Orb – Sensing | -1.871 | 0.256 |
|  | Orb – Sheet | -1.318 | 0.294 |
|  | Orb – Space | -1.699 | 0.136 |
|  | Orb – Specialist | -1.481 | 0.350 |
|  | Ambush – Ground | -0.708 | 0.704 |
|  | Ambush – Other | -0.677 | 0.727 |
|  | Ambush – Sensing | -0.639 | 1.309 |
|  | Ambush – Sheet | **0.010** | **1.457** |
|  | Ambush – Space | -0.560 | 1.246 |
|  | Ambush – Specialist | -0.242 | 1.458 |
|  | Ground – Other | -0.311 | 0.456 |
|  | Ground – Sensing | -0.498 | 1.281 |
|  | Ground – Sheet | **0.243** | **1.113** |
|  | Ground – Space | -0.305 | 1.080 |
|  | Ground – Specialist | -0.005 | 1.346 |
|  | Other – Sensing | -0.682 | 1.125 |
|  | Other – Sheet | **0.164** | **1.002** |
|  | Other – Space | -0.406 | 0.946 |
|  | Other – Specialist | -0.145 | 1.263 |
|  | Sensing – Sheet | -0.529 | 1.230 |
|  | Sensing – Space | -1.002 | 1.058 |
|  | Sensing – Specialist | -0.628 | 1.289 |
|  | Sheet – Space | -1.020 | 0.329 |
|  | Sheet – Specialist | -0.660 | 0.735 |
|  | Space – Specialist | -0.532 | 1.219 |

Cases where the 95% confidence interval does not overlap zero, and therefore provides evidence of a significant effect, are highlighted in bold
